# Approximating complex musculoskeletal biomechanics using multidimensional autogenerating polynomials

**DOI:** 10.1101/759878

**Authors:** Anton Sobinov, Matthew Boots, Valeriya Gritsenko, Lee E. Fisher, Robert A. Gaunt, Sergiy Yakovenko

## Abstract

Computational models of the musculoskeletal system are scientific tools used to study human movement, quantify the effects of injury and disease, and plan surgical interventions. Additionally, these models could also be used to intuitively link biological control signals and realistic high-dimensional articulated prosthetic limbs. However, implementing fast and accurate musculoskeletal computations that can be used to control a prosthetic limb in real-time is a challenging problem. As muscles typically span multiple joints, the wrapping over complex geometrical constraints changes their moment arms and length as a function of joint angle and, thus, their ability to generate joint torques. As a result of these biomechanical complexities, calculating these muscle state variables in real-time is a difficult simulation problem. Here, we report a method to accurately and efficiently calculate these variables for the forearm muscles that actuate the hand and wrist across multiple postures. The posture dependent muscle geometry, moment arms and lengths of modeled muscles, were captured using autogenerating polynomials that expanded their optimal selection of terms using information measurements. The iterative process approximated 33 musculotendon actuators, each spanning up to 6 DOFs in an 18 DOF model of the human arm and hand, defined over the full physiological range of motion. Using these polynomials, the entire forearm anatomy could be computed in <10 µs, which is far better than what is required for real-time performance, and with low errors in moment arms (below 5%) and lengths (below 0.4%). Moreover, we demonstrate that the number of elements in these autogenerating polynomials does not increase exponentially with the increase in complexity of muscles, increasing linearly instead. The similar structure and function of muscles are represented with specific invariant polynomial terms. Dimensionality reduction using the polynomial terms alone resulted in clusters comprised of muscles with similar functions, suggesting that the polynomials themselves captured biologically relevant features of muscle structure and function. We propose that this novel method of describing musculoskeletal biomechanics might further improve the applications of detailed and scalable models for the description of human movement.

## Introduction

The remarkable dexterity of the hand results from the coordinated motion of 27 kinematic degrees of freedom (DOF) actuated by arm and hand muscles. This problem of coordination is solved continuously by our neuromuscular system without perceived cognitive effort. Yet, for prosthetic applications, the current approaches, such as pattern recognition and mode switching require significant training time (Cordella et al., 2016). Moreover, the skill and cognitive load required for continuous prosthetic control increases with the number of available prosthetic DOFs (Deeny et al., 2014). This phenomenon is captured by *the dimensionality curse* problem in movement planning, which occurs due to the increasing volume of possible solutions with the increasing number of dimensions. Recently, machine learning statistical methods have gained popularity in computer vision and robotic control problems of comparable complexity. In particular, deep learning algorithms are capable of remarkable performance in vision and language tasks (Riesenhuber and Poggio, 1999) and significantly outperform the shallow networks that had been common for decades. These performance gains and the resistance to the dimensionality curse are enabled by the hierarchical processing inherent in these multilayer deep networks, which is a biomimetic property common to biological cortical networks (Poggio et al., 2017). However, training these deep networks requires large amounts of labelled data and usually results in a black-box transformation, without any transparent internal mechanisms that would generate insights into the underlying control scheme (reviewed in Lapuschkin et al., 2019). In addition, machine learning solutions often require episodic model retraining (Hermann et al., 2015), and rely on a considerable memory space to store the necessary parameters (Weston et al., 2014). These constraints pose significant challenges for real-time control systems for both phenomenological and mechanistic models of human hand biomechanics. Overall, this approach limits our understanding of model boundaries, the reliable domain of operation, and, importantly, the principles of the modelled system that can be tested and improved further. Instead, using mechanistic alternatives based on known biology may overcome these limitations.

Transforming biological signals into intended prosthetic movements using biomimetic principles may solve the problem of integration between the biological and technological control systems. These systems may often be at odds with each other due to the discord in expected and executed movement. Thus, the challenges of biomimetic approaches are in specifying and implementing valid motor control theories. One such dominant theory focuses on internal models expressed within the nervous system (Angelaki et al., 2004; Kawato, 1999; Wolpert et al., 1998); it embodies an engineering concept termed the Smith predictor (Smith, 1957). This theory relies on accurate estimates of the controlled plant to overcome both nonlinear dynamics and temporal delays. Another complimentary concept is neuromechanical tuning (Prochazka and Yakovenko, 2007; Sreenivasa et al., 2019; Ting et al., 2007), which deals with the nature of computed signals within the closed-loop control system and postulates reliance on the coupled interplay between neural and mechanical dynamics. The key idea of these theories is that body dynamics and musculoskeletal (MS) biomechanics are essential components that require valid models (Ting et al., 2015; Ting and Chiel, 2017) or good-enough biomimetic approximations within the design of a robotic prosthesis (Kumar et al., 2013). The recent use of MS models for human-machine interfaces (Crouch and Huang, 2016) shows promising results for this type of approach.

MS modelling is an important scientific tool in theoretical motor control (Berniker et al., 2009; Lillicrap and Scott, 2013; Winter, 2009) and its applications in human-machine interfaces (Crouch and Huang, 2016; Thorsen et al., 2001). MS models are typically comprised of geometrical descriptions of each joint’s degrees of freedom (DOFs) and muscle paths around these DOFs. A muscle’s contribution to joint torque depends on the distance to the DOF axis of rotation, called moment arm, and muscle state described by its length and velocity that alter force generation (An et al., 1984; Zajac, 1989). Calculating these musculoskeletal kinematic variables in a specific posture requires computation of the shortest path between the points of attachment in the presence of objects like bones and other muscles around which a muscle wraps (Delp et al., 2007). Software packages like OpenSim (SimTK) provide tools for computation of kinematic variables based on a 3D model of a limb or whole body. These calculations are very computationally costly and can only be performed in real-time for simple models. However, models of increasing complexity are required in both research and applications, rapidly raising their computational cost to burdensome levels.

The computational load of MS models has led to the development of multiple approximation methods that improve computational efficiency. Menegaldo and colleagues (Menegaldo et al., 2004) proposed a series of multidimensional polynomials describing the MS variables of human leg muscles. Later these polynomials were used to simulate the musculotendon dynamics of upper (Rankin and Neptune, 2012) and lower limbs (Chadwick et al., 2009). This approach supports very high computational performance with low requirements on the available memory and the number of mathematical operations. However, the generalizability of this method is limited by the hand-selected polynomial structure, which begins to have significant errors in the more complex biomechanical scenarios that occur in the hand. Addressing this limitation is not trivial. Defining the polynomial structure itself becomes considerably more difficult as the biomechanical complexity of the musculoskeletal system increases. For example, muscles actuating the thumb may cross seven DOFs (three wrist and four thumb), potentially resulting in a 7-dimensional polynomial to describe its behavior. Another approach developed by Sartori and colleagues (Sartori et al., 2012) emphasizes the quality of approximation using cubic splines. Albeit being computationally expensive, the ability of this approach to operate at real-time has been shown in a 3-DOF per muscle model (Durandau et al., 2018). The drawback of cubic splines, however, are their limited scalability: the number of spline coefficients increases exponentially with the number of DOFs that the muscle crosses. Ultimately, both methods aim to simplify complex musculoskeletal calculations and exhibit problems with accommodating the increasing model complexity, severely limiting the possibility of MS structure analysis and application.

In this study, we present an information theory-based algorithm of polynomial approximation of MS kinematic variables that scales linearly with the complexity of the model. The resolution of the dimensionality problem in the approximations is addressed with the search for the optimal structure of approximating functions. We assess the quality in terms of approximation error and evaluation time on a MS model with 33 musculotendon actuators crossing multiple DOFs each (up to 6 DOFs per muscle). Our mathematical model belongs to the class of phenomenological descriptions capturing the input-output relationship and, yet, it may also represent theoretical principles associated with muscle function as has been demonstrated for several models of this type (Frigg and Hartmann, 2018). Thus, the structure of the produced optimal polynomials is analyzed not only in terms of muscle anatomy but also its function.

## Methods

The approximation of muscle path kinematic variables consisted of three steps: *i*) creating a dataset describing muscle length and moment arm values for all physiological postures using the OpenSim model; *ii*) searching for a set of optimal polynomials approximating kinematic variables implemented with a physical constraint between muscle moment arms and muscle length; and *iii*) validating the produced polynomials.

### Dataset

We used a previously developed model of the arm and hand to capture the relationship between muscle lengths and moment arms in all physiological postures (Boots et al., 2017; Gritsenko et al., 2016). The model contains 33 musculotendon actuators, some representing multiple heads of the same muscle, spanning 18 physiological DOFs (see Tables 1 and 2 in Appendix) and was implemented in OpenSim software (Delp et al., 2007). Similar to the previous study of Sartori et al. (Sartori et al., 2012) the values for the kinematic variables were obtained on a uniform grid with 9 points per DOF, resulting in the domain size of 9^*d*^ data points per muscle, where *d* is the number of DOFs that a muscle crosses. The extreme positions were included so that 9 points were selected within the range from 0% to 100% of DOF range. For example, since the *extensor carpi ulnaris* muscle spans two DOFs (wrist flexion-extension and pronation-supination) in our model (ulna deviation is not simulated) its moment arms and muscle lengths were sampled in 9^2^=81 positions. This 9-point dataset contained 674,937 points. In addition, to compare the approximations achieved with different methods (described below), we generated the 8-point dataset containing 348,136 values that fits between data in the 9-point dataset.

**Table 1.** Polynomial term notation and kinematic *muscle invariants*. (*x*_1_, *x*_2_, *x*_3_, *x*_4_) are coordinates.

| $v$-axis index | Unique combination | Examples of polynomial terms | $K$-notation |
| --- | --- | --- | --- |
| 1 | (1) | $x_1, x_2$ | $K_1, K_2$ |
| 2 | (2) | $x_1^2$ | $K_{11}$ |
| 3 | (3) | $x_1^3$ | $K_{111}$ |
| 4 | (4) | $x_1^4$ | $K_{1111}$ |
| 5 | (5) | $x_1^5$ | $K_{11111}$ |
| 6 | (1,1) | $x_1x_2, x_2x_3$ | $K_{12}, K_{23}$ |
| 7 | (1,2) | $x_1^2x_2, x_2x_3^2$ | $K_{112}, K_{233}$ |
| 8 | (1,3) | $x_1x_2^3$ | $K_{1222}$ |
| 9 | (1,4) | $x_1x_2^4$ | $K_{12222}$ |
| 10 | (2,2) | $x_1^2x_2^2$ | $K_{1122}$ |
| 11 | (2,3) | $x_1^2x_2^3$ | $K_{11222}$ |
| 12 | (1,1,1) | $x_1x_2x_3$ | $K_{123}$ |
| 13 | (1,1,2) | $x_1x_2x_3^2$ | $K_{1233}$ |
| 14 | (1,1,3) | $x_1x_2x_3^3$ | $K_{12333}$ |
| 15 | (1,2,2) | $x_1x_2^2x_3^2$ | $K_{12233}$ |
| 16 | (1,1,1,1) | $x_1x_2x_3x_4$ | $K_{1234}$ |
| 17 | (1,1,1,2) | $x_1x_2x_3x_4^2$ | $K_{12344}$ |
| 18 | (1,1,1,1,1) | $x_1x_2x_3x_4x_5$ | $K_{12345}$ |

**Table 2.** Model performance comparison. Cubic spline (CS) and two polynomial approximations with and without the constraint linking muscle lengths and moment arms (constrained and unconstrained polynomials, CP and UP), as described by algorithm in Model Physical Constraints in Methods. L is length, MA is moment arms.

| Method | RMS error ± standard deviation, % | Total number of parameters | AIC, au |
| --- | --- | --- | --- |

|  | L | MA | L | MA | L | MA |
| --- | --- | --- | --- | --- | --- | --- |
| CS | $1.34*10^{-5} \pm 1.56*10^{-5}$ | $1.84*10^{-6} \pm 2.47*10^{-6}$ | $1.1*10^9$ | $1.64*10^{10}$ | $2.2*10^9$ | $3.2*10^{10}$ |
| UP | $0.0383 \pm 0.0918$ | $0.757 \pm 1.477$ | 610 | 705 | $-6.7*10^6$ | $-5.7*10^5$ |
| CP | $0.0382 \pm 0.0910$ | $0.757 \pm 1.477$ | 661 | 783 | $-6.7*10^6$ | $-5.7*10^5$ |

### Model Structure

Moment arms and muscle lengths were approximated with a polynomial described by Eq. 1.

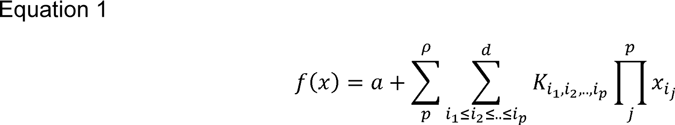

where *a* is an intercept, *ρ* is the selected maximum of polynomial power, *d* is the number of DOFs, *x* =(*x*_1_,..,*x*_*d*_) is the state vector with values of angles at each DOF, *K* is the multidimensional matrix of polynomial term coefficients, sum and product coefficients (*p, i*, and *j*) iterate from 1. The **polynomial structure** is then defined by the non-zero values of *K* and a parameters. For example, *extensor carpi ulnaris* moment arms (with *ρ* = 4, *d* = 2) were described by the polynomial structures (*a, K*_1_,*K*_2_, *K*_11_, *K*_12_, *K*_22_,*K*_111_, *K*_112_, *K*_122_, *K*_222_, *K*_1111_, *K*_1112_, *K*_1122_, *K*_1222_) around elbow extension-flexion (e-f) and (*a, K*_1_, *K*_2_,*K*_11_,*K*_12_,*K*_22_, *K*_111_, *K*_112_,*K*_122_, *K*_222_, *K*_1111_, *K*_1112_, *K*_1122_, *K*_1222_, *K*_2222_) around wrist supinatison-pronation (s-p) (Fig. 2B), where indices 1 and 2 correspond to wrist pronation-supination and flexion-extension, respectively.

**Figure 1.**
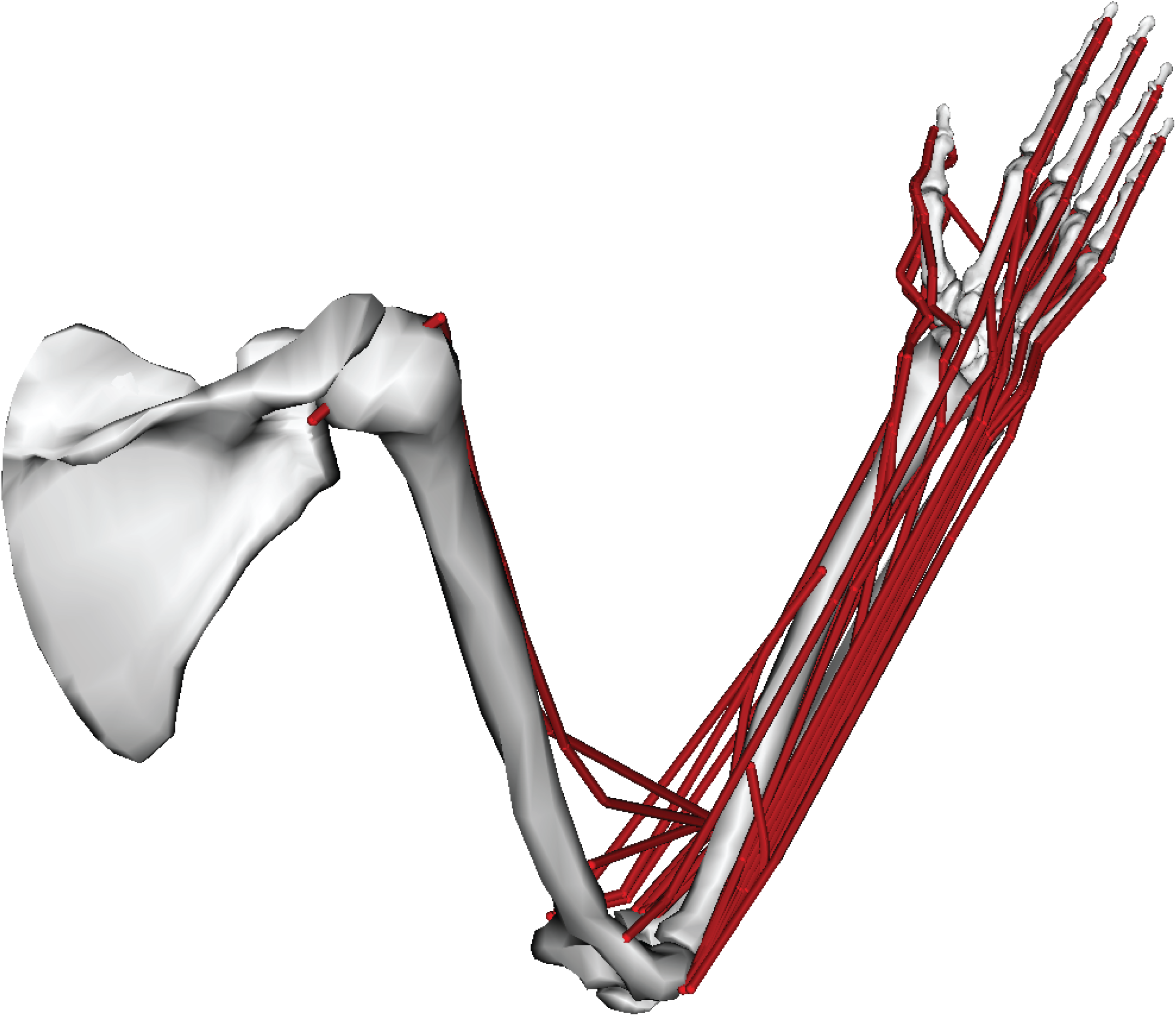
Upper-limb representation in OpenSim. The geometry of muscle paths is shown in red for the displayed posture.

**Figure 2.**
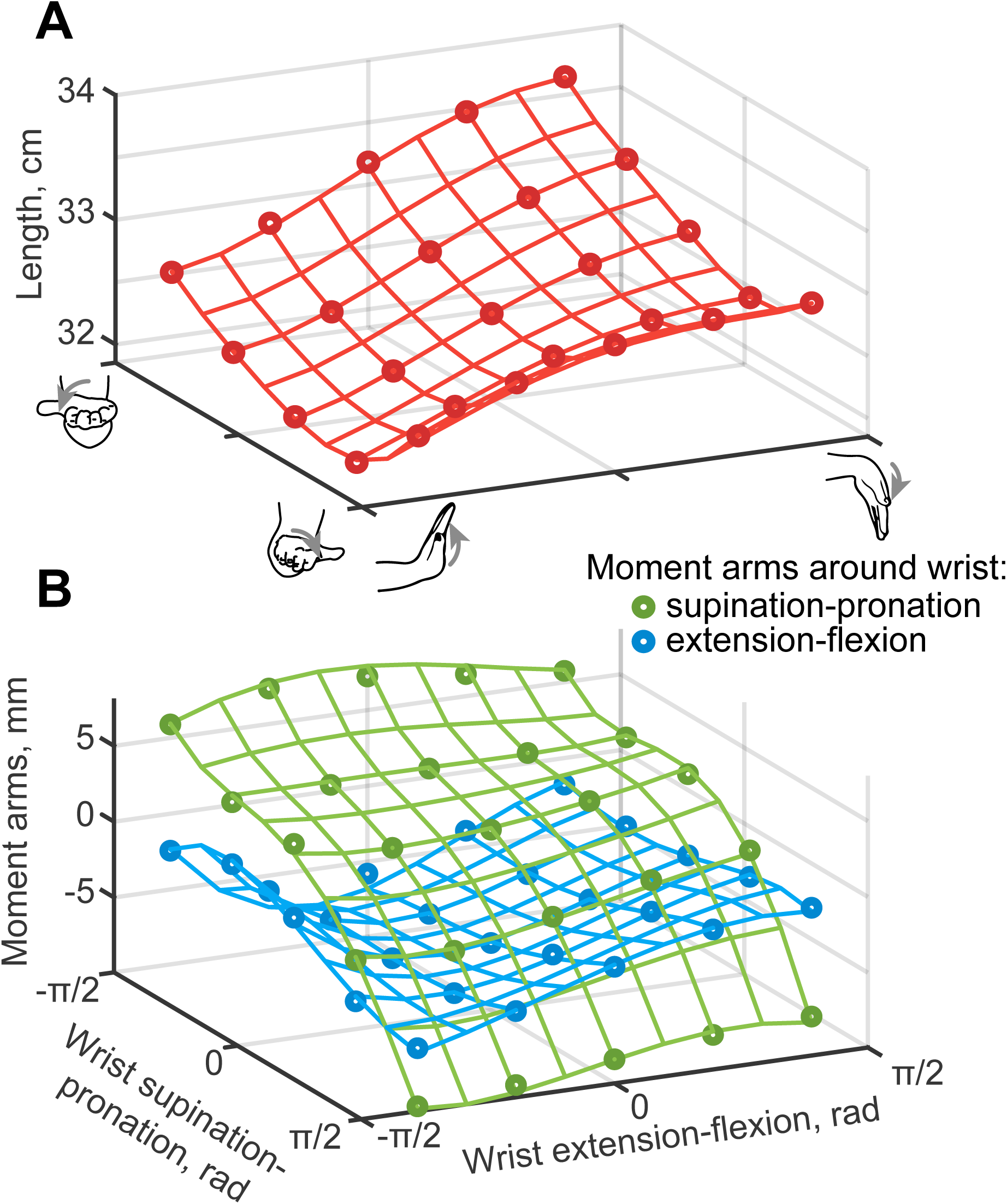
The example of kinematic approximation for *extensor carpi ulnaris* muscle. **A**. The muscle path length is shown as a function of wrist e-f and s-p DOFs, with points from OpenSim model fitted with the continuous functions plotted as a wireframe. **B**. The two corresponding moment arm relationships are shown for the same domain of postures.

### Model Physical Constraints

Moment arms can be estimated as a partial differential of the muscle length in local coordinates (An et al., 1984; Brand et al., 1975):

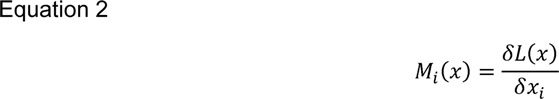

where *i* is the index of a DOF actuated by the muscle, *x*_*i*_ is the coordinate of *i*th DOF, *M*_*i*_ (*x*) is the posture-dependent function of the moment arm around *i*th DOF, *L*(*x*) is the muscle length function. The kinematic variables of a given muscle are then captured by a single function *L*(*x*) and a set of functions {*M*_*i*_ (*x*)} for muscles spanning multiple DOFs.

The following algorithm finds a new function *L*(*x*) and updates its set of moment arm functions {*M*_*i*_} in agreement with the relationship in Eq.2:

1. Calculate a set of intermediate muscle length polynomials 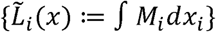.
2. Combine the terms of *L*(*x*) and 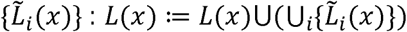.
3. Differentiate analytically the polynomial L(x) (Eq.2) to update the complimentary set of moment arm functions, {*M*_*i*_ (*x*)}.
4. Calculate *a* and *K* coefficients in *L*(*x*) and {*M*_*i*_ (*x*)} using the original dataset.

For example, for an arbitrary muscle spanning two DOFs *x*= (*x*_1_, *x*_2_) with its length described by a function 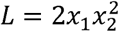, we have a polynomial term *x*_1_*x*_2_*x*_2_, which is denoted by the term *K*_122_. Similarly, the corresponding two moment arm functions 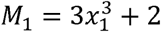 and *M*_2_ = 5*x*_1_*x*_2_ are described by the terms (*K*_111_,*a*) and (*K*_12_). The integrals of *M*_1_, *M*_2_ in step 1 are: 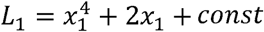 or structure 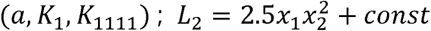 or structure (*a, K*_122_). In step 2, the ensemble function *L*(*x*) adhering to Eq. 2 will be 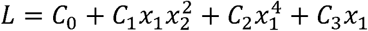 where *C*_*i*_ are scalar coefficients in the structure (*a,K*_1_, *K*_122_, *K*_1111_). This step embeds the differential relationship between path length and its moment arms. In step 3, the moment arms are 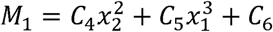 or structure (*a, K*_22_,*K*_111_) and *M*_2_ = *C*_7_*x*_1_*x*_2_ or structure (*K*_12_). We introduce this additional notation for constants to separate them from polynomial structures. We used a linear pseudoinverse on the original dataset to calculate the coefficients *C*_0-7_. These coefficients were used to evaluate the quality of fit (next section) and to analyze the nature of embedded information within the polynomials (see below, Kinematic Muscle Invariants).

### Model generation and validation

The geometries of muscle wrapping around joints vary greatly in their complexity and, consequently, their model representations. The simplest muscles can be approximated with a constant if their path is posture independent, and complex muscles may involve many polynomial terms. The search for the optimal model requires the evaluation of each additional term from the domain of terms that grows exponentially with the number of actuated DOFs. Thus, muscles crossing 6 DOFs in our model were the most challenging. To solve this, we created an optimization algorithm similar to forward stepwise regression (Izenman, 2008). This method was adapted to include all possible polynomial terms and the constraint in Eq.2 in the process of expanding the polynomial structure with additional terms until the information tradeoff indicated overfitting. For this purpose, we used the corrected Akaike Information Criterion (AICc) for a finite sample size (Akaike, 1974; Burnham and Anderson, 2004):

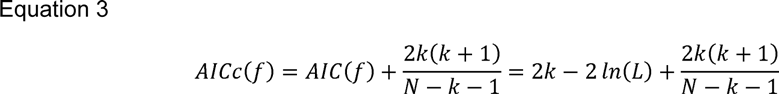

where *f* is an approximation function, is the Akaike Information Criterion, *k* is the number of parameters in the model, *N* is the number of data points, and *L* is a maximum likelihood estimation of the polynomial representing this dataset. The peak value of *L* for the normally distributed estimated residuals is *ln*(*L*) = −0.5*N*(*ln*(2*π σ*^2^+ 1) = -*N ln*(*σ*) + *const*, where *σ ln*(*L*) is the root-mean-square (RMS) error. The model-independent constants are ignored in the substitution of in Eq.3 because we use AICc values to compare multiple models (see further details on pp. 62-67 in Burnham and Anderson, 2004):

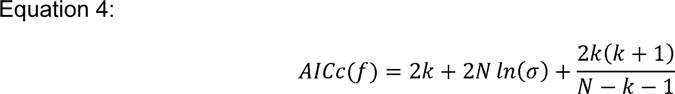

To remove potential differences between DOFs, we normalized the muscle length values to the range of motion and the moment arm values to their maximum across all physiological postures.

The analysis selected the terms of the polynomial structure for a muscle as follows (Fig. 3A):

**Figure 3.**
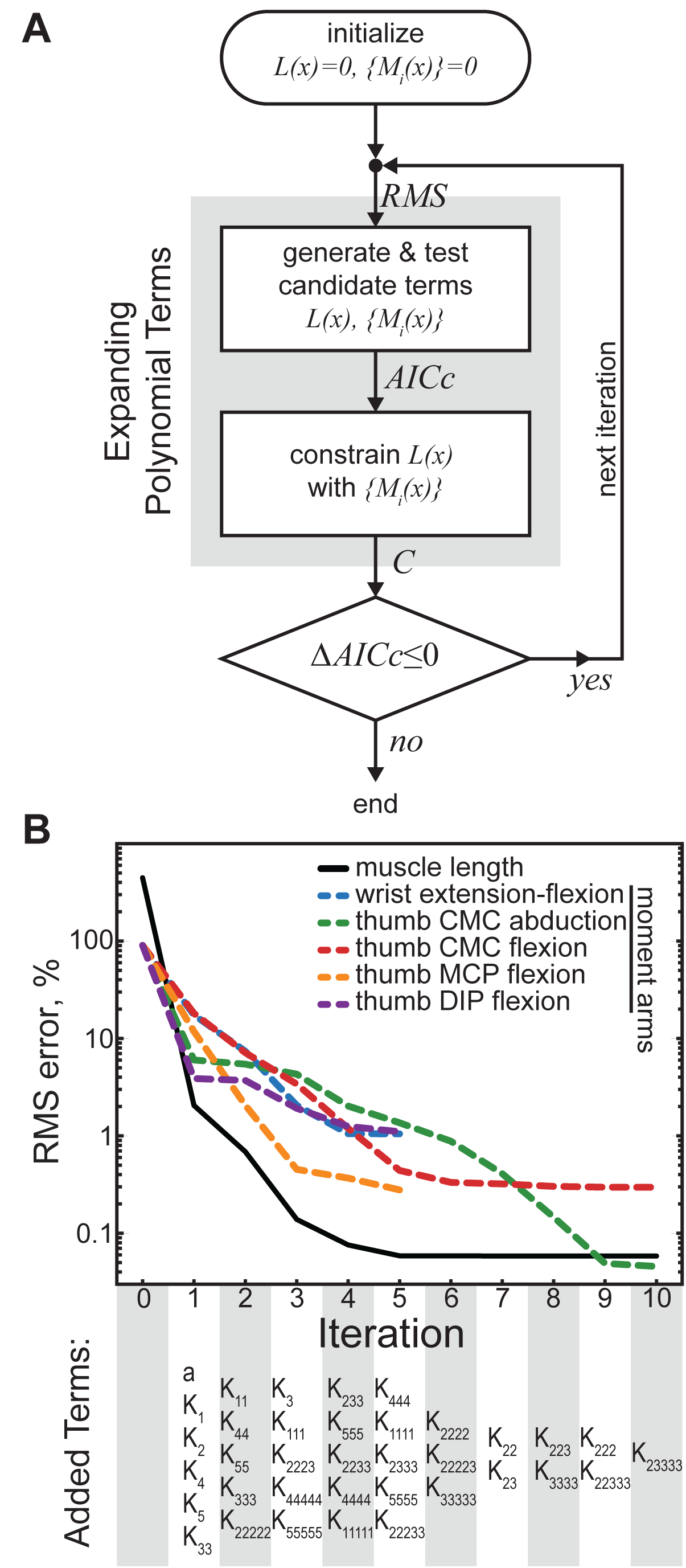
Optimal model generation algorithm. **A**. The optimization flow schematic showing the flow of calculations using the amalgamated algorithm of model generation with physical constraint. RMS errors of model performance are computed at the onset of each new iteration and followed by the expansion of polynomial candidates. The process continues while there are improvements in AICc metric. **B**. Example of generating the system of polynomial functions describing *flexor pollicis longus*. The decrease in RMS errors for all DOFs actuated by this muscle were plotted for each iteration of the algorithm. The progression of terms added to minimize AICc in 6 polynomials is shown below the plot.

1. Initialize a variable (empty polynomial without terms) for the functions approximating muscle length *L*(*x*) and its set of moment arm functions, {*M*_*i*_ (*x*)}.
2. Make a list of potential candidates for the expansion of each polynomial using all possible combinations from the fifth degree polynomial: *ψ* (*L*); {*ψ*(*M*_*i*_)}_*i*_.
3. Select optimal functions indicated by the smallest AICc values from the lists *ψ*(•) and append them to the current approximation: 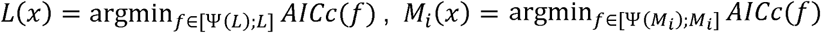.
4. Use the algorithm, described above (Model Physical Constraints), to impose the relationship of Eq. 2.
5. Return to step 2: i) if further expansion is possible (*ψ*(*L*) or *ψ*(*M*_*i*_) are not empty), and ii) the change in AICc values is negative between iterations.

The progression of model assembly with this algorithm can be seen in Fig. 3B showing the optimization of kinematic variables for *flexor pollicis longus* with the iterative expansion. The first evaluation of errors was performed relative to zero model (*L*(*x*) = 0;{*M*_*i*_ (*x*)} = 0). The errors for the selected terms were evaluated in the following iteration step. In the first iteration, the muscle length was approximated by (*a, K*_1_,*K*_2_, *K*_4_,*K*_5_, *K*_33_), where some terms came from the selection of terms in step 3 and the rest from the integration in step 4. In the second iteration, the approximation was expanded using elements *K*_11_, *K*_44_,*K*_55_, *K*_333_, *K*_2222_, and the precision of muscle length fit decreases below 1%. In the fifth iteration, only thumb carpometacarpal (CMC) & metacarpophalangeal (MCP) moment arms required further optimization when other DOFs reached the minimum of AICc. In the tenth iteration, the evaluation of optimal parameter selection was finished with the high precision of 10^−3^ for the fit of muscle length across all physiological postures. Here, the worst moment arm fit of wrist extension-flexion (dashed blue line) was 1.05% in units normalized to the range of motion and the maximum magnitude of moment arm or 0.2 mm in absolute units.

The accuracy of polynomial fit generally increases with the number of terms in the polynomial structure. For each iteration, the selection of potential candidates for expansion, Ψ(*P*(*x*)), contains polynomials with all terms of *P*(*x*) and one additional term from the possible additional terms in a polynomial of the same power. For example, let *P*(*x*) be a two-dimensional polynomial with structure (*a, K*_1_,*K*_11_), full 2-dimensional polynomial of power 2 has a structure (*a, K*_1_,*K*_2_, *K*_11_, *K*_12_, *K*_22_). Then the list of potential candidates is: *Ψ*(P(*x*))= [(*a,K*_1_, *K*_2_,*K*_11_); (*a,K*_1_, *K*_11_, *K*_12_); (*a, K*_1_,*K*_11_,*K*_*22*_)]. The size of Ψ(*P*(*x*)) increases when higher power terms are required.

### Similarity index

Muscles with similar function may require similar approximation structures to capture their kinematics. To test this idea, we used a measure of similarity between polynomial structures. Consider polynomials *L*_*A*_ and *L*_*B*_ characterizing muscles A and B. Each polynomial can be described by a collection of shared or common terms (*P*_*C*_) and a collection of non-common terms (*P*_*NC*_), so that *L*_*A*_ = *P*_*C*_ ∪ *P*_*ANC*_ and *L*_*B*_ = *P*_*C*_ ∪ *P*_*BNC*_, where *P*_*ANC*_ are the terms present in *L*_*A*_ and not in *L*_*B*_ and *P*_*BNC*_ are the terms present in *L*_*B*_ and not in *L*_*A*_. Then, the similarity index (SI) is calculated as:

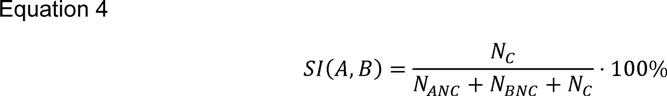

where *N*_*C*_, *N*_*ANC*_, *N*_*BNC*_ are the number of terms in *P*_*C*_, *P*_*ANC*_, *P*_*BNC*_, respectively. *SI* equals to 100% when two polynomials have completely identical structures (*K* terms), and to 0% when they are completely different.

### Kinematic Muscle Invariant

Additional details describing polynomial composition was captured using muscle representation in a Euclidean space formed by the basis of unique polynomial power terms (*K, Table 1*). Here, the obvious similarity due to mechanical actions around the same DOFs was removed (using *v*-axis index, Table 1) to test if the approximations contained additional functional relationships. Whether or not functional information is embedded in the pattern of polynomials could then be tested by examining the distance between muscles in this space. For the full polynomial of power *ρ* = 5 and maximum muscle dimensionality *d* = 6 these unique combinations are the following: [(1, 1, 1, 1, 1), (1, 1, 1, 1), (1, 1, 1, 2), (1, 1, 1), (1, 1, 2), (1, 1, 3), (1, 1), (1, 2, 2), (1, 2), (1, 3), (1, 4), (1), (2, 2), (2, 3), (2), (3), (4), (5)], where (1, 1, 1, 1, 1) is, e.g., *x*_1_*x*_2_ *x*_3_*x*_4_*x*_5_ and (5) is 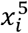. The coefficients for these ordered 18 combinations defined the coordinates of a vector representing a given muscle-length polynomial. We converted all polynomials into unit vectors with the normalized sums of coefficients of the same terms from different DOFs, 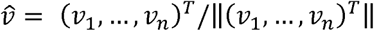. For example, for 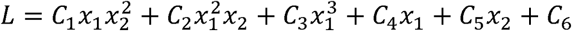, the vector has nonzero elements [*v*_9_ = |*C*_1_| + | *C* _2_| *v*_12_ = | *C* _4_| + |*C*_5_| *v*_16_ = |*C*_3_|]. Structural difference of two polynomials can then be obtained as a distance between their vectors. We call vectors of each muscle in the basis of v*-*axes as *muscle invariants*. The structural difference between muscles is minimal when power composition of all terms and their absolute coefficients are similar in both polynomials even if they cross different DOFs, and large when their power compositions do not have the same terms.

### Memory and Time

The computer memory required for spline approximation was calculated as a size of MATLAB’s ‘.mat’ files that contained single-precision spline parameters saved using ‘-v7.3’ flag which enables compression. The computer memory required for polynomials was calculated as the size of executable ‘.mexw64’ files compiled with Visual Studio 2017 C++ with ‘/O2’ optimization. Evaluation time was obtained using MATLAB’s Profiler. Individual samples for mean and standard deviation of evaluation time were obtained per muscle’s dataset during estimation of fit quality. All computations were done on DELL Precision Workstation T5810 XL (Intel Xeon processor E5-2620 v3 2.4 GHz, 64 GB DDR4 RAM, SK Hynix SH920 512 GB SSD) running Windows 10.

### Statistics

The accuracy of polynomials was analyzed with standard statistical tools. The RMS error values were used to evaluate errors in the approximated values relative to the dataset used for fitting and the independent testing dataset (see above, Dataset). We detected outliers using a method similar to (Sartori et al., 2012), which resulted in the removal of less than 0.09% of values from the 9-point dataset. We estimated maximum expected error using Chebyshev’s theorem with 1% significance level. Linear regression was used to test the relationship between the complexity of functions represented by the number of DOFs a muscle spans and the complexity of the approximating polynomials.

The similarity of muscle invariants 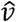 across multiple muscle groups was tested with dimensionality reduction analyses, i.e. principle component analysis (PCA) and hierarchical clustering. The Euclidean distance between vectors was first analyzed with the average linkage hierarchical clustering implemented in SciPy. Then, the dominant relationships in this distribution of *muscle invariants* were analyzed with PCA (Arisman, 2014; Scikit-learn module in Pedregosa et al., 2011).

The representation of structural and functional information within the muscle length invariants was further tested by comparing the distributions of the distances between muscle pairs with similar structure or similar function to muscles with different structure or different function. These distributions were shown to be non-normal using D’Agostino’s K-squared test (D’Agostino and Pearson, 1973) that measures deviation from the normal skewness and kurtosis. We used one-tailed Mann-Whitney *U* test (Mann and Whitney, 1947) to assess the two hypotheses that functional and structural similarities are represented in the colocalization of the *muscle invariants*. In general, this test was used to assess the likelihood of observing a smaller distance between the randomly selected pairs of *muscle invariants* with matching function or structure than the distance between the randomly selected pairs with shuffled function or structure. The smaller distances between the pairs in matched populations than the larger distances between the pairs from the shuffled populations were also tested with one-sided sign test (Conover, 1999). The symmetrical distribution of samples around the mean is not assumed in the sign test; thus, it is a better choice for this problem then Wilcoxon signed-rank test. All tests were performed with the conservative value of α set at 0.01.

## Results

We developed a precise and efficient method to describe the MS kinematics of a human forearm and hand, extending previous work with approximation functions (Menegaldo et al., 2004; Sartori et al., 2012). Here, we formalized the dynamic selection of terms in a best-fit polynomial function using a quantitative tracking of overfitting. Moreover, we used the differential relationship between muscle length and moment arms within the derivation algorithm to generate mutually consistent analytical models of these two variables. We tested if the composition of polynomials embedded information about muscle structure and/or function.

### Approximation of muscle lengths and moment arms

We subdivided values in the dataset (see above) into two groups for creating models and their testing. All best-fit models, splines and polynomials, approximated moment arms with <5% error and muscle length with <0.4% error, as shown in Fig. 4 and Table 2.

**Figure 4.**
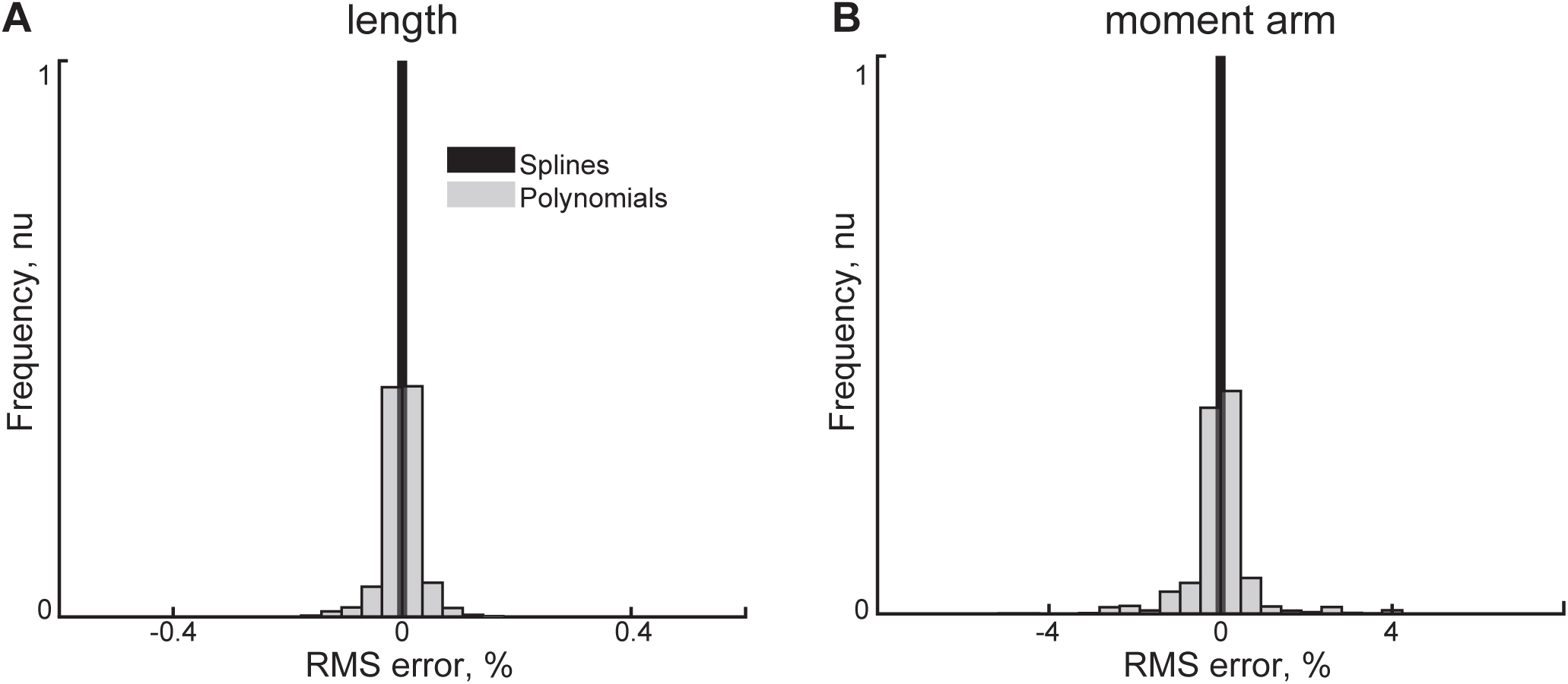
The distributions of normalized errors in the estimation of muscle lengths (**A**) and moment arms (**B**) are shown for two models (splines and polynomials). The histogram frequency was normalized to the total count of samples.

Although the approximation error with splines was the lowest, the implementation of splines required the highest number of parameters – eight orders of magnitude difference (compare cubic splines and constrained polynomials in Table 1). The large number of parameters in the cubic spline model exceeded the number of values in the dataset, which corresponded to impractical AICc values. We used AIC values instead to compare the relative quality of models: the constrained polynomial values were −6.7*10^6^ and −5.7*10^5^, as compared to the cubic spline values 2.2*10^9^ and 3.2*10^10^. This difference indicates the preference of AIC metric to the constrained polynomial model. The addition of model physical constraints (Eq. 2) to the polynomial generation algorithm did not significantly change the precision of the polynomial model (p>0.9) with similar errors and AIC values in Table 2. The histograms of error distributions were superimposed in Fig. 4. The length approximation errors in Fig. 4A were smaller than those of moment arm errors in Fig. 4B, as expected from Eq.2. In general, the differentiation process increases the magnitudes of errors.

A small portion of values in the datasets were marked as outliers and removed from further analyses: unconstrained polynomials had 0.08% muscle length outliers and 0.03% moment arm outliers; constrained polynomials had 0.08% and 0.03%, respectively. No spline errors were marked as outliers.

Both polynomial models were over 7000 times faster than the cubic spline (Table 3) and required 2.8*10^5^ times less memory. The search time for the constrained polynomials was 3.3 times faster than that for the unconstrained polynomials with the increase in performance gained when the selection of polynomial terms originated in the relationship between muscle length and moment arms.

**Table 3.**
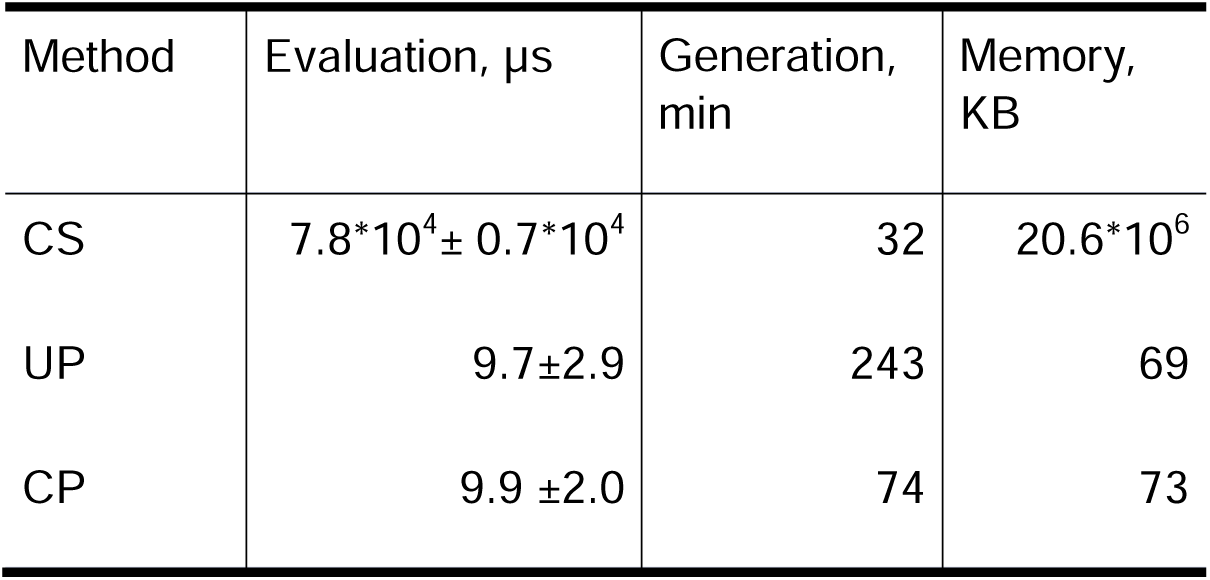
Time and memory requirements of approximations methods for kinematic variables.

### Structure of Approximating Polynomials

Both the constrained and unconstrained polynomial models were similar in composition as determined by the high similarity between the two models (Fig. 5A). Because the constrained muscle length function has higher polynomial power than its moment arm functions, we used *ρ* = 4 to generate Ψ(*M*_*i*_), and *ρ* = 5 to generate Ψ(*L*). The similarity index is high when both models contain the same polynomial terms, which is indicated by the predominance of high similarity indices for all muscles in Fig. 5A. It takes about 20 terms per muscle to achieve high accuracy (Fig. 5B). The average similarity between muscles was 87.1%, and the biggest difference was observed in three muscles *biceps brachii short head, flexor carpi radialis*, and *adductor pollicis transversus* with similarity indices at about 60%. This indicates that the compositions of constrained and unconstrained polynomial models were similar.

**Figure 5.**
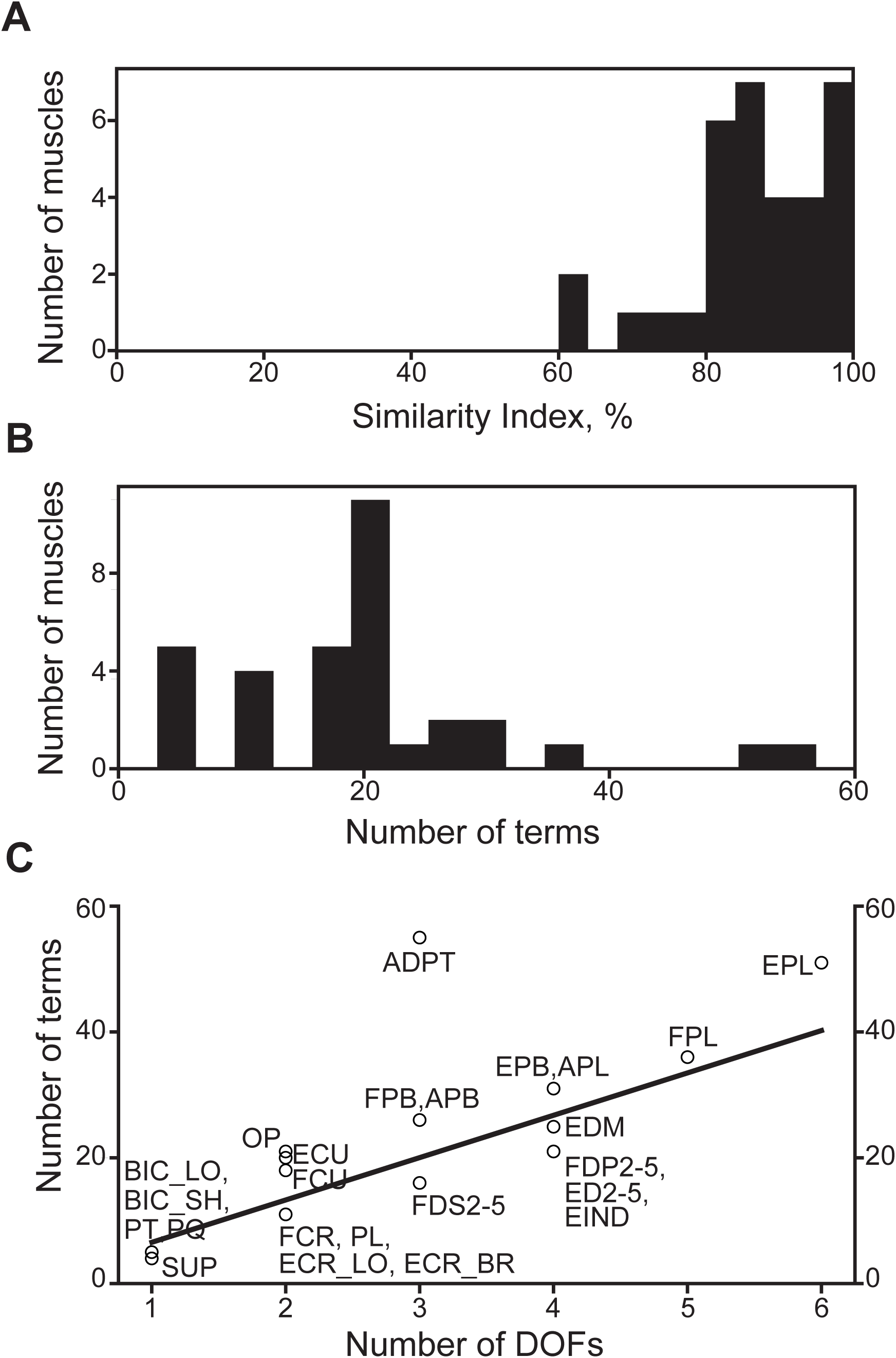
Complexity of muscle structures. **A.** Similarity index between functions approximating muscle lengths generated with and without the physical constraint imposed by Eq. 2 in step 4 of the above algorithm. **B.** The distribution of polynomial complexity expressed as the number of terms. **C**. The relationship between the number of terms in the muscle length polynomial (circles) and the number of DOFs the muscle spans (line, *y* = 6.73*x* – 0.16, *r* = 0.74, *p* < 2 · 10^−6^).

The increase in anatomical complexity indicated by the number of DOFs actuated by a muscle was predicted to correspond to the exponential increase in the number of terms required. This type of relationship was evident in the cubic spline model, where thumb muscles spanning up to 6 DOFs required the highest number of parameters. It is remarkable that the relationship between the number of terms in the muscle length polynomial and the number of DOFs the muscle spans is instead linear (*r* = 0.74, Fig. 5C). Moreover, the model fractional complexity, measured as the ratio of terms selected to all possible terms available, decreased as the number of DOFs controlled by a muscle increased (Supplementary Fig. 1, *r* = −0.83). The most complex muscles in our model were the thumb muscles (ADPT, FPB, APB, EPB, APL, FPL, EPL), and they appeared above the regression line (Fig. 5C). Instead, the finger muscles (FDS2-5, FDP2-5, ED2-5, EDM, EIND) were below the line (Fig. 5C), suggesting that these muscles have a lower relative complexity than the thumb muscles.

### Structure and Function

We hypothesized that the generated models capture structural and functional features of muscles and developed a measure of embedded muscle attributes, coined *muscle invariants*. These muscle invariants represent each muscle in the space of polynomial term powers. To avoid trivial relationships where similarity could be simply determined by the index of DOF actuated by a pair of muscles, we removed DOF identity information and preserved only the power signature of each term. The difference between muscles was captured as Euclidean distances between their vectors. To visualize the 18-dimensional space of all power terms (Table 1), the distance heatmap was calculated between all muscle pairs (Fig. 6A), and the corresponding vectors were plotted in the axes of two main principle components computed with PCA (Fig. 6B). The clustering algorithm generated the dendrogram based on these distances. A selection of distal thumb muscles (ADPT, APB, OP, APL) was visibly separated from about 6 other subgroups, with the closest subgroup formed by another subset of thumb muscles (EPL and EPB). These groups were separated by the dashed line in the dendrogram of Fig. 6A. The thumb muscles were followed (top to bottom) by: *extensor carpi radialis* and wrist flexors (ECR_LO, ECR_BR, FCR, PL), *flexor pollicis brevis* and *extensor carpi ulnaris* (FPB and ECU), finger and wrist flexors and extensors, wrist rotators located in the forearm (FDP2-4, FDS3-5, ED2, ED4, ED5, EIND, PL, FCR, PQ, PT, SUP), the rest of digit muscles with *flexor carpi ulnaris* (ED3, EDM, FDS2, FDP5, FCU, FPL), and *biceps brachii* (BIC_SH, BIC_LO).

**Figure 6.**
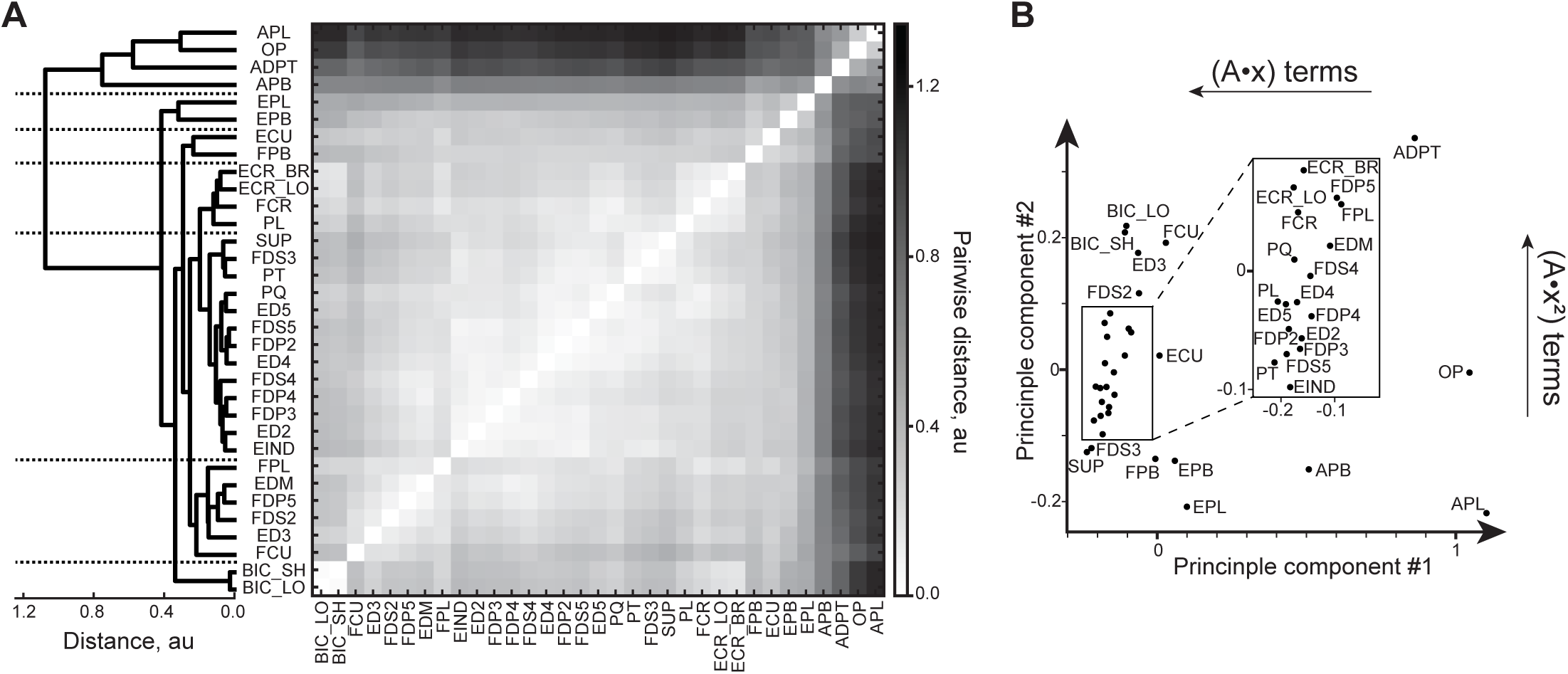
Kinematic muscle invariants. **A**. Average-linkage dendrogram computed from the heatmap of pairwise distances between muscle invariants. Horizontal dashed lines indicate subgroups described in text. **B**. The representation of muscle invariants in the space of their main two principle components. *Insert*: expanded view of a portion of the plot.

The differences between muscle invariants were largely captured by the first two principal components (86% of variance explained). Their largest coefficients were associated with linear 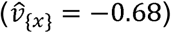 and square 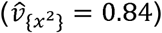 powers of polynomial terms. The linear relationship between joint angle and muscle length corresponds to a semi-circle muscle path around a joint. This simplistic behavior is characteristic for 1-DOF finger joints, muscles in the bottom-left corner and the insert of Fig. 6B. Muscles in the bottom-right corner, e.g., thumb muscles, used less linear terms than other muscles. Overall, the space of muscle invariants has a nonrandom and hierarchically structured pattern.

We tested if muscle invariants contain information about their anatomical location by comparing Euclidian distances between the invariants with shared DOFs. Since there is a limited set of muscles that do not span the same joints, we tested the idea that those pairs of muscles that share a given DOF would be closer to each other than those that do not share that DOF. We assigned phalangeal DOFs (MCP, PIP, DIP) to be different to each other, but the same across fingers 2-5 because of their similarity and the lack of intrinsic hand muscles (e.g., lumbricals) in the model. This selection ensured local structural similarity in the group with a shared DOF (Fig. 7A, blue) and local difference in the group without a shared DOF (Fig. 7A, red), but it did not prevent the selection of muscle pairs in each group based on their structure relative to other DOFs. Fig. 7A shows the probability of observing a given distance between a pair of muscles with a shared DOF and without a shared DOF based on 1306 and 1862 pairs, respectively. The selection of muscles into these groups was executed sequentially by examining all muscles for each DOF in the model. The difference distribution between the two distributions in Fig.7A shown in Fig. 7B was computed by examining the difference between each pair with a shared DOF and comparing it with each pair that had one of the two muscles in the group without a shared DOF, resulting in 20,746 comparisons.

**Figure 7.**
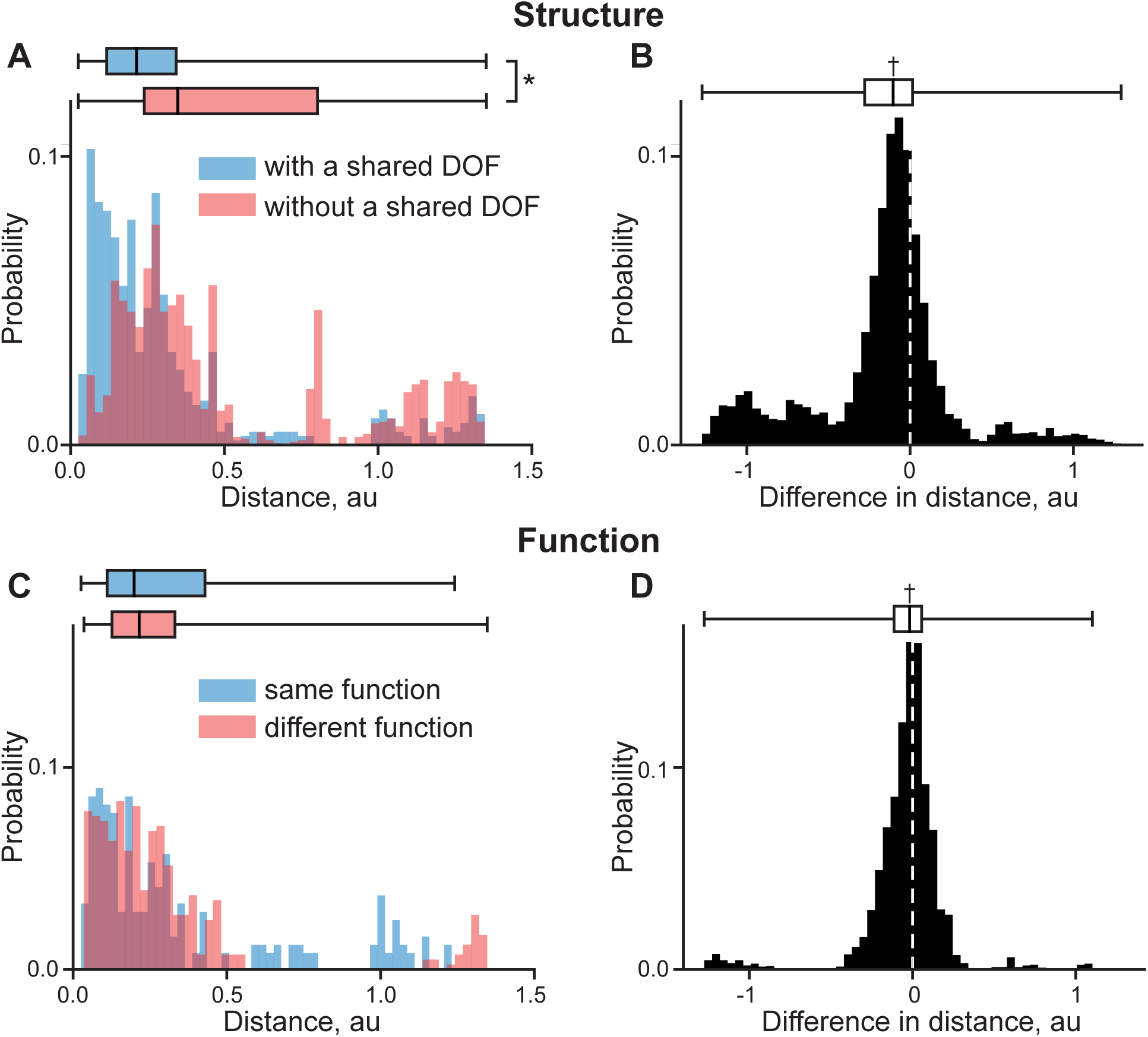
The structural and functional information embedded in muscle invariants. **A**. The probability distributions of observing the distance between the pairs of muscle invariants with (blue) and without (red) a shared DOF. **B**. The test of difference between the two groups. **C**. The probability distributions of the distance between the pairs with the shared structural information and with (blue) and without (red) shared functions. **D**. The test of difference between the two groups. Box plots indicate a median and 25^th^-75^th^ quantile region. The significant differences between the overlap of distributions tested with Mann-Whitney U test is marked with (*). The sign test significance is marked with (†).

The median of difference was significantly different from zero (−0.10, sign test *p* < 10^−8^). Both groups were not normally distributed (D’Agostino’s K-squared test of normality, *p* < 10^−8^) and similar anatomical pairs were closer to each other which was evident from the non-equal distribution of the two groups (Mann-Whitney U test: *U*= 7 · 10^5^, p< 10^−8^). We found that the muscle invariants capture the structural information related to the identity of their actuated DOFs.

We tested if the muscle invariants contain functional information beyond that explained by the anatomical similarities. For this purpose, we defined seven functional categories based on their primary mechanical function: wrist supinators (BIC_LO, BIC_SH, SUP), pronators (PT, PQ), extensors (ECR_LO, ECR_BR, ECU), flexors (FCR, FCU, PL), finger flexors (FDS2-5, FDP2-5), extensors (ED2-5, EDM, EIND), and thumb muscles (APL, OP, APB, EPL, EPB, FPB, FPL, ADPT). We tested the idea that two muscles from the same category are closer together than those from different categories even when all these muscles actuate the same DOF. For this reason, we selected all pairs of muscles with (490 pairs) and without (816 pairs) a shared function selected from the seven categories and computed the distance between these pairs, shown in Fig. 7C. Next, we computed the distance between the two groups based on the combinations of all these pairs (3496 samples), shown in Fig. 7D. The distributions in Fig. 7C were also not normal (*p* < 10^−8^). While distributions of the two groups were overlapping (*p* = 0.61), the median difference between them was significantly different from zero (−0.02, sign test *p* < 10^−8^). This supports the hypothesis that DOF-independent functional differences are captured by the muscle invariants.

## Discussion

We approximated MS kinematics of the human forearm and hand with a new type of autogenerating model that embeds biomechanical constraints between muscle parameters. The model reached optimal performance with polynomial simulations showing high precision and computational efficiency. While the model was developed as a descriptive tool, the fine details captured within the muscle-posture relationships include the differential connection between moment arms and muscle lengths and reflect the high-level mechanistic properties of forearm and hand muscle function. The composition of terms in these models was objectively determined by the embedded information and demonstrated the patterns associated with anatomy and function. The mechanical specification of muscles for the control of different hand DOFs and different functions has not been previously demonstrated.

All models are simplifications or approximations of reality, but some approximations are useful. The complex geometric interactions—sliding and wrapping—between muscles and other mechanical body structures pose a considerable computational challenge for real-time applications (Blana et al., 2017). The engineering trade-off between complexity, performance, and accuracy pushed the development of simplified biomechanical limb models that assumed constant moment arm and posture relationships (Crouch and Huang, 2016) or reduced the span of musculotendon anatomy to ease computational demand (Durandau et al., 2018). Here, we report a method of capturing the kinematic MS transformations within the biomechanical model of the forearm and hand that does not require these simplifications. The approximating models can be mechanistic or phenomenological. The goal of phenomenological model is to capture the input-output relationship without the effort of describing the mechanistic explanation present within this transformation. We show that our objective method of generating the approximations captured structural and functional features of MS organization in the phenomenological model.

### Autogenerating models

Interest in MS approximations has been steadily increasing with the development of computational tools for human motion analysis, e.g., OpenSim (Delp et al., 2007). Accuracy of these approximations has been demonstrated with B-spline models (Durandau et al., 2018; Sartori et al., 2012) and computational efficiency has been achieved with polynomial models (Chadwick et al., 2009; Menegaldo et al., 2004). The optimal polynomials derived in this manuscript have the benefits of both accuracy and computational efficiency.

The manual subjective selection of polynomial terms for each muscle is usually based on the number of DOFs the muscle crosses, the quality of simulation, and the numerical cost of evaluating functions. In contrast, our optimization algorithm chooses the polynomial terms objectively based on the information criterion to reflect objective dependencies within the data. The information criterion is a type of cost function that allows comparison between different polynomial models and prevents overfitting with an excessive number of terms. The latter is possible when using the subjective desired precision of fit, as in (Chadwick et al., 2009). Similar to (Menegaldo et al., 2004) the number of terms in the optimized polynomial grows with the number of muscle’s DOFs, but the term composition varies to reflect the diverse anatomy and function.

We found multiple levels of structure embedded in the power composition of polynomial terms. A linear relationship between muscle length and joint angle is characteristic for 1-DOF finger joints. The near-linear relationship between moment arm profile and joint angle we showed in thumb muscles has been commonly observed in other studies (Loren et al., 1996; Menegaldo et al., 2004). The physiological function of this relationship could be associated with compensation for the muscle force-length relationship at the edges of the range of motion. The diverse function and behavior of thumb muscles found during movement (Kaufman et al., 1999) is mirrored in our results by their separation from other muscles and high variability between each other.

Previously we have examined the grouping of muscles based on their length-posture relationships where the similarity between muscles was determined by common muscle length shortening and lengthening in response to postural changes (see Fig.7 in Gritsenko et al., 2016). The current analysis of muscle organization does not separate antagonistic muscles, with the focus only on the polynomial sets that shape muscle paths. Similar to the previous analysis, thumb muscles are clearly separated from other finger muscles. We have also included muscles with antagonistic functions in separate groups in the analysis of muscle properties captured by the polynomials (Fig. 7C,D). This test indicated a functional difference between the muscle invariants even when the differences accounted for by muscle location were removed; albeit, this difference was small. This result supports the idea that the commonly observed muscle synergies during movement can be at least in part explained by the structure and function embedded in the musculotendon paths.

### Real-time high-dimensional musculoskeletal computations

The optimal polynomials efficiently compute highly complex MS kinematics for real-time applications. The polynomials describing 33 musculotendon actuators each crossing up to 6 DOFs can be evaluated within 10 µs, requiring less than 75 KB of RAM. To contrast, the previous state-of-the-art performance for a lower-limb model with 13 musculotendon actuators, each crossing up to 3 DOF was shown to be less than 2.5 ms (Durandau et al., 2018). Our more than hundred-fold time efficiency improvement on the method was also accompanied by a similar improvement in required memory (about 10MB worth of coefficients in (Durandau et al., 2018) based on (Sartori et al., 2012)). The improvements are largely due to the exponential rise in the required computational resources with the dimensionality increase of the spline model, as previously shown (Sartori et al., 2012) and by our implementation. This ‘dimensionality curse’ may prevent the application of splines in complex models recently developed for offline analyses (Holzbaur et al., 2005; Paclet and Quaine, 2012; Rajagopal et al., 2016). Our optimal polynomial approach shows linear scaling of the model (Fig. 5C) allowing these models to be used in real-time applications.

The described optimization algorithm is structurally similar to stepwise regression (Izenman, 2008), but has several important differences. First, it automatically constructs and explores all possible polynomial combinations of the input variables within reasonable power limitations. Second, our algorithm uses AIC (Akaike, 1974; Burnham and Anderson, 2002) instead of F-statistic as the objective measure of improvements. The additional term in AIC takes into account the trade-off between the quality of fit and the increased model complexity. This is a novel use of information measures (Akaike, Bayesian and other) that have been previously used mostly as a stopping criterion (Bendel and Afifi, 1977). An information criterion allows flexibility when choosing the tradeoff between quality of fit and the measure of model complexity. For example, using the number of processor commands instead of the number of variables for each term is useful for the development of extremely high-performing routines or for computationally costly devices, like portable chips or graphics processing units. Third, our approximation algorithm embeds the differential relationship between muscle length and its moment arms in the search for the best polynomial coefficients. This novel approach of using the formulation of structural constraints within the algorithm decreased model assembly time. These approximations are ready to be used on a portable device that requires a real-time simulation of MS variables, e.g., a biomimetic prosthesis or a medical assessment device.

### Limitations

We chose to implement the fitting algorithm with the use of polynomial sequences as the most accurate representation of the MS relationships. The alternative implementations could use sequences of trigonometric or exponential terms. For example, any data with periodic relationships would be efficiently represented by trigonometric functions, and any data with sigmoidal transitions or limits of range could be represented by exponential functions. However, the relationships between moment arms and posture are smooth because of soft tissue properties. In this case, we can rely on the theoretical conclusion from Taylor’s theorem stating that any smooth function can be described with a polynomial approximation. Then the only potential failures would be the observations of discontinuities in the muscle properties. We have indeed observed sharp transitions which are always associated with geometric model failures where a muscle path slipped off a wrapping surface. These behaviors were detected and corrected prior to the approximation (Boots et al., 2017). Thus, our model is appropriate for the physical system it represents.

The autogenerating polynomials were iteratively created with the selection of a single term per equation at a time. This enabled fast optimization of the full system of equations describing moment arms and muscle lengths. It is possible that multiple terms can be more optimal than a single term. This would be indicated by the premature termination of the optimization routine even when a more optimal solution is available for multiple terms selected in the same iteration. We tested this eventuality by repeating the model generation with an algorithm capable of adding one or two terms per iteration per equation. This method produced the same solutions for our dataset (data not shown), but the evaluation time increased by an order of magnitude as compared to the standard method.

The current method is limited to the description of forearm muscles in a generic representation of the human hand. Future analysis of validated models that span the shoulder will improve our understanding of muscle specialization. We expect to see new functional groups with the structure different from that of any of the hand functional groups because of the unique biomechanics of the shoulder joint (Lucas, 1973; Voisin, 2006). These functional groups can be then further refined by their evaluation on models with subject-specific segment scaling and morphometric differences (Akita and Nimura, 2016). It will be also illuminating to compare the muscle organization of the upper limb to that of the lower limb, considering their proposed coevolution (Rolian et al., 2010), covariability in developmental modules (Hallgrímsson et al., 2002) and high observed topological similarity (Diogo et al., 2013) in humans. However, accurate and valid lower-limb models are still under development.

### Conclusions

We approximated the kinematic variables for human hand and forearm muscles with both high precision (<5% error across 18 DOFs) and efficiency (<75 KB, <10 μs). Adding the differential relationship between moment arms and muscle lengths improved solutions and the speed of their calculations. The approach overcomes the curse of dimensionality and scales linearly with increased complexity for large MS models. The structural content of optimal polynomials reflects muscle anatomy and function. This novel description can be further applied in neuromechanics and its applications.

## Supplementary information

The validity of selecting the sampling rate of the relationship between posture and muscle parameters was tested by comparing the quality of approximation with three different rates, i.e., the training datasets were sampled at 3, 5, and 9 values per degree of freedom (DOF). The corresponding three testing datasets with data points residing between the training data points were used for validation. The overall fitting errors were not significantly different between 5- and 9-point datasets. However, infrequent failures in the 5-point model were effectively resolved with the 9-point model. It is likely that further increases in the sampling rate are not likely to increase model performance and may lead to the overfitting by exceeding the quality of the musculoskeletal representation in OpenSim. Since the 5-point model had a very similar performance to the 9-point model, it can be effectively used as an intermediate fast approximation for iterative adjustments needed to validate muscle geometry against experimental data (as in Boots et al., 2017). Overall, the 9-point model was deemed to be optimal.

**Supplementary Table 1.** The list of simulated DOFs. Each label describes both a DOF and the direction of axis using the following structure: <LIMB>_<JOINT>_<MIN>_<MAX>, where LIMB corresponds to the limb where the joint is located, i.e. ‘ra’ stands for ‘right arm’, JOINT is the joint of this DOF, e.g., ‘wr’ is ‘wrist’. Digit joints have their identifying number: 1 thumb; 2 index; 3 middle; 4 ring; and, 5 pinky. The last two suffixes MIN and MAX indicate the anatomical direction of axis, e.g., ‘ra_wr_s_p’ indicates the range of the wrist pronation-supination DOF (−1.5708 rad for the supinated posture and the maximum 1.5708 rad for the pronated posture).

| DOF ID | Label | Range | Description of action |
| --- | --- | --- | --- |

|  |  | [MIN, MAX], rad |  |
| --- | --- | --- | --- |
| 1 | ra_wr_s_p | -1.5708, 1.5708 | wrist pronation/supination motion |
| 2 | ra_wr_e_f | -1.2217, 1.2217 | wrist flexion/extension motion |
| 3 | ra_cmc1_f_e | 0, 0.8727 | thumb proximal flexion/extension motion |
| 4 | ra_cmc1_ad_ab | 0, 0.8727 | thumb proximal abduction/adduction motion |
| 5 | ra_mcp1_f_e | -0.7854, 0 | thumb central flexion/extension motion |
| 6 | ra_ip1_f_e | -1.5708, 0 | thumb distal flexion/extension motion |
| 7 | ra_mcp2_e_f | 0, 1.5708 | index proximal flexion/extension motion |
| 8 | ra_pip2_e_f | 0, 2.0944 | index central flexion/extension motion |
| 9 | ra_dip2_e_f | 0, 1.5708 | index distal flexion/extension motion |
| 10 | ra_mcp3_e_f | 0, 1.5708 | middle proximal flexion/extension motion |
| 11 | ra_pip3_e_f | 0, 2.0944 | middle central flexion/extension motion |
| 12 | ra_dip3_e_f | 0, 1.5708 | middle distal flexion/extension motion |
| 13 | ra_mcp4_e_f | 0, 1.5708 | ring proximal flexion/extension motion |
| 14 | ra_pip4_e_f | 0, 2.0944 | ring central flexion/extension motion |
| 15 | ra_dip4_e_f | 0, 1.5708 | ring distal flexion/extension motion |
| 16 | ra_mcp5_e_f | 0, 1.5708 | pinky proximal flexion/extension motion |
| 17 | ra_pip5_e_f | 0, 2.0944 | pinky central flexion/extension motion |
| 18 | ra_dip5_e_f | 0, 1.5708 | pinky distal flexion/extension motion |

The list of simulated DOFs. The last two suffixes separated by underscores indicate the direction of the motion, e.g., “\_s\_p” indicates the motion from supinated to pronated posture.

**Supplementary Table 2.** The list of simulated musculotendon actuators. Brief labels used in figures are shown with their anatomical names and the corresponding information about number and identity of actuated DOFs, as described in Supplementary Table 1.

| Muscle ID | Label | Musculotendon actuator | Number of DOFs | DOF IDs |
| --- | --- | --- | --- | --- |
| 1 | BIC_LO | *Biceps brachii long head* | 1 | 1 |
| 2 | BIC_SH | *Biceps brachii short head* | 1 | 1 |
| 3 | SUP | *Supinator* | 1 | 1 |
| 4 | PT | *Pronator teres* | 1 | 1 |
| 5 | PQ | *Pronator quadratus* | 1 | 1 |
| 6 | ECR_LO | *Extensor carpi radialis longus* | 2 | 1 2 |
| 7 | ECR_BR | *Extensor carpi radialis brevis* | 2 | 1 2 |
| 8 | ECU | *Extensor carpi ulnaris* | 2 | 1 2 |
| 9 | FCR | *Flexor carpi radialis* | 2 | 1 2 |
| 10 | FCU | *Flexor carpi ulnaris* | 2 | 1 2 |
| 11 | PL | *Palmaris longus* | 2 | 1 2 |
| 12 | FDS5 | *Flexor digitorum superficialis (pinky finger)* | 3 | 2 16 17 |
| 13 | FDS4 | *Flexor digitorum superficialis (ring finger)* | 3 | 2 13 14 |
| 14 | FDS3 | *Flexor digitorum superficialis (middle finger)* | 3 | 2 10 11 |
| 15 | FDS2 | *Flexor digitorum superficialis (index finger)* | 3 | 2 7 8 |
| 16 | FDP5 | *Flexor digitorum profundus (pinky finger)* | 4 | 2 16 17 18 |
| 17 | FDP4 | *Flexor digitorum profundus (ring finger)* | 4 | 2 13 14 15 |
| 18 | FDP3 | *Flexor digitorum profundus (middle finger)* | 4 | 2 10 11 12 |
| 19 | FDP2 | *Flexor digitorum profundus (index finger)* | 4 | 2 7 8 9 |
| 20 | EDM | *Extensor digiti minimi* | 4 | 2 16 17 18 |
| 21 | ED5 | *Extensor digitorum (pinky finger)* | 4 | 2 16 17 18 |
| 22 | ED4 | *Extensor digitorum (ring finger)* | 4 | 2 13 14 15 |
| 23 | ED3 | *Extensor digitorum (middle finger)* | 4 | 2 10 11 12 |
| 24 | ED2 | *Extensor digitorum (index finger)* | 4 | 2 7 8 9 |
| 25 | EIND | *Extensor indicis* | 4 | 2 7 8 9 |
| 26 | EPL | *Extensor pollicis longus* | 5 | 1 2 4 3 5 6 |
| 27 | EPB | *Extensor pollicis brevis* | 4 | 2 4 3 5 |
| 28 | FPB | *Flexor pollicis brevis* | 3 | 4 3 5 |
| 29 | FPL | *Flexor pollicis longus* | 5 | 2 4 3 5 6 |
| 30 | APL | *Abductor pollicis longus* | 4 | 1 2 4 3 |
| 31 | OP | *Opponens pollicis* | 2 | 4 3 |
| 32 | APB | *Abductor pollicis brevis* | 3 | 4 3 5 |
| 33 | ADPT | *Adductor pollicis transversus* | 3 | 4 3 5 |

## Acknowledgements

We thank Jennifer Collinger for critical comments on the manuscript. Research was sponsored by the U.S. Army Research Office and the Defense Advanced Research Projects Agency (DARPA) was accomplished under Cooperative Agreement Number W911NF-15-2-0016. The views and conclusions contained in this document are those of the authors and should not be interpreted as representing the official policies, either expressed or implied, of the Army Research Office, Army Research Laboratory, or the U.S. Government. The U.S. Government is authorized to reproduce and distribute reprints for Government purposes notwithstanding any copyright notation hereon.

